# c-Myc plays a key role in IFN-γ induced persistence of *Chlamydia trachomatis*

**DOI:** 10.1101/2021.03.09.433696

**Authors:** Nadine Vollmuth, Sudha Janaki-Raman, Lisa Schlicker, Naziia Kurmasheva, Werner Schmitz, Almut Schulze, Thomas Rudel, Karthika Rajeeve

## Abstract

*Chlamydia trachomatis (Ctr)* can persist over long periods of time within their host cell and thereby establish chronic infections. One of the major inducers of chlamydial persistence is interferon-gamma (IFN-γ) released by immune cells as a mechanism of immune defence. IFN-γ activates the catabolic depletion of L-tryptophan (Trp) via indoleamine 2,3-dioxygenase (IDO), resulting in persistent *Chlamydia*. Here we show that IFN-γ depletes c-Myc, the key regulator of host cell metabolism, in a STAT1-dependent manner. Expression of c-Myc rescued *Chlamydia* from IFN-γ-induced persistence in cultured cell lines, but also in human fallopian tube organoids. L-tryptophan concentrations control c-Myc levels via the PI3K-GSK3ß axis. Unbiased metabolic analysis revealed that *Chlamydia* infection reprograms the host cell tricarboxylic acid (TCA) cycle to support pyrimidine biosynthesis. Addition of TCA cycle intermediates or pyrimidine/purine nucleosides to infected cells rescued *Chlamydia* from IFN-γ-induced persistence. Thus, our results challenge the longstanding hypothesis of L-tryptophan depletion through IDO as the major mechanism of IFN-γ-induced metabolic immune defence and significantly extends the understanding of the role IFN-γ as a broad modulator of host cell metabolism.

## Introduction

*Chlamydia trachomatis* (*Ctr*) is an obligate intracellular human pathogen, which causes a broad range of acute and chronic diseases^1^. It is the leading cause of bacterial sexually transmitted diseases (STD), with more than 130 million new cases annually^2^. Infection of the urogenital tract by *Chlamydia* can lead to urethritis, infertility, elevated risk of foetal and neonatal deaths, ectopic pregnancies and pelvic inflammatory disease (PID)^3,4^. Furthermore, left untreated, *Chlamydia* infection increases the risk of HIV infections^5,6^ and might contribute to the development of cervical and ovarian cancer^7–10^. The pathogen extensively interferes with the physiology of the infected cell, since it depends entirely on its host cell as a replicative niche during infection^11^.

*Chlamydiae* have a unique biphasic life cycle, consisting of two physiologically and morphologically distinct forms^12^. This gram-negative bacterium initiates its developmental cycle by attachment and invasion of the host cell by the elementary body (EB), the non-dividing infectious form of the pathogen. Inside the cell, EB stay within the endosomes, which they modify rapidly to create a replicative niche termed ‘inclusion’, to avoid lysosomal degradation^13^. EB differentiate into reticulate bodies (RB), which is the non-infectious replicating form of the pathogen. It takes several rounds of cell division until RB re-differentiate back into EB and the progenies are released by host cell lysis or extrusion, an exocytosis like mechanism, to infect neighbouring cells^14,15^. Apart from active infection, *Chlamydia* can turn into a dormant state called persistence and can remain over a long time, maybe even years^16^ within its host cell and thereby establish a chronic infection. During the persistent state, the pathogen remains viable and replicates its genome, but it exhibits decreased metabolic activity and inhibited cell division, which leads to the formation of enlarged pleomorphic aberrant bodies (AB)^17^. The transition to the persistent form might represent an important chlamydial survival mechanism against antibiotics and the immune response of the host, since this process is reversible^17,18^. The conversion to this dormant stage is induced by penicillin^19^, iron deficiency^20^, amino acid starvation^21^ and interferon-gamma (IFN-γ)^17,18^. IFN-γ is an immune regulated cytokine, which is involved in the cell-intrinsic immunity against several intracellular pathogens, including *Chlamydia^22^*. The downstream signalling pathways activated by IFN-γ are highly species specific and differ dramatically between mouse and human cells^23^. In human cells, the anti-chlamydial effect of IFN-γ is predominantly mediated by the induction of indoleamine-2,3-dioxygenase (IDO), an enzyme that catalyses the initial step of L-tryptophan (Trp) degradation to N-formyl kynurenine and kynurenine^24^. Trp is a crucial amino acid for chlamydial development^25,26^, and its depletion leads to persistence^18,27^.

IFN-γ belongs to the type II interferons that bind to the extracellular domain of the interferon-gamma receptor, which is a heterodimer of the two subunits IFNGR1 and IFNGR2. The intracellular domains of the IFNGR1 subunits are associated with Janus kinase 1 (Jak1), while the IFNGR2 subunits are associated with Jak2. Activation of Jak1 and Jak2 results in phosphorylation of the receptor and subsequent recruitment and phosphorylation of signal transducer and activator of transcription (STAT1). STAT1 phosphorylation at tyrosine 701 and serine 727 leads to its homodimerization and nuclear translocation. Once in the nucleus, STAT1 homodimers bind to IFN-γ-activated sequence (GAS) elements in the promoters of target genes to regulate their transcription^28–30^. IFN-γ can function as both, a growth inhibiting or promoting cytokine, in a STAT1-dependent manner^31,32^. Binding of STAT1 homodimers to the consensus GAS elements in the *c-myc* promoter inhibits its expression transcriptionally^28^. Concurrently, *c-MYC* expression is not only negatively regulated by STAT1 but *stat1* is also a negative target gene of c-Myc. Hence, c-Myc and STAT1 regulate each other in a negative feedback loop at the transcriptional level^33^.

The transcription factor c-Myc targets genes involved in the regulation of numerous cellular processes, such as cell proliferation, cell growth, translation, metabolism and apoptosis^34–37^. Furthermore, c-Myc activity increases energy production, anabolic metabolism, promotion of aerobic glycolysis and glutaminolysis, inducing mitochondrial biogenesis and tricarboxylic acid (TCA) cycle activity^37^. Glutaminolysis increases the production of biomass by providing TCA cycle intermediates via anaplerosis, which enhances their availability for the production of amino acids, nucleotides and lipids^38^. c-Myc also critically controls nucleotide biosynthesis by directly regulating the expression of genes that encode the enzymes involved in the production of precursors of all nucleotides^39,40^. For example, c-Myc directly controls the cis-regulatory element in the 5’-UTR of phosphoribosyl pyrophosphate synthetase (PRPS), which catalyses the first committed step in purine biosynthesis^41^. In pyrimidine biosynthesis, the rate-limiting step is catalysed by carbamoyl aspartate dehydratase (CAD)^42^, which is also regulated by c-Myc as a response to growth stimulatory signals, such as activation of EGFR/RAS/MAP kinase^43^, Hif 1/2 α^44^ or estrogen receptor/Sp1^45^ pathways. We recently identified a central role of c-Myc in the control of the metabolism in *Chlamydia-infected* cells^46^. *Chlamydia* has a reduced genome and only very limited metabolic capacity. For example, they have a truncated TCA cycle and lack the ability to synthesize purine and pyrimidine nucleotides *de novo* but acquire ATP and nucleosides from the host cell^47,48^.

Expression of IDO and depletion of Trp has been previously described as the main reason for interferon-induced persistence in *Chlamydia*^49,50^. Here we show that IFN-γ induces a STAT1-dependent depletion of c-Myc in *Chlamydia-infected* cells, which de-regulates the host metabolism and induces *Chlamydia* persistence. Importantly, addition of the TCA cycle intermediate α-ketoglutarate or pyrimidine/purine nucleosides was sufficient to prevent persistence and restore chlamydial replication. Our data demonstrate a central role of c-Myc-regulated metabolic pathways in the IFN-γ-induced persistence of *C. trachomatis*.

## Results

### IFN-γ treatment prevents c-Myc induction and impairs the development of *Chlamydia*

c-Myc is an important regulator of host cell metabolism and is indispensable for chlamydial acute infection and progeny formation^46^. Since *Chlamydia* obtain all nutrients from the host cell and it has been shown that several environmental conditions affecting host cell metabolism like iron and amino acid starvation also induce chlamydial persistence^51,52^, we investigated the role of c-Myc in chlamydial persistence. From our previous study, it was known that ablation of c-Myc expression interfered with chlamydial development and progeny formation^46^. To investigate if *Chlamydia* enter a persistence state in c-Myc depleted cells, we infected a HeLa 229 cell line with an anhydrotetracycline (AHT)-inducible short hairpin RNA (shRNA) for c-Myc in the absence and presence of the inducer. In agreement with our previous results^46^, silencing of c-Myc expression prevented inclusion formation (Fig. 1a). We then cultivated cells for 24 h with AHT to suppress c-Myc expression and then either added AHT for another 12 h or removed AHT to re-establish c-Myc expression (see scheme Fig. 1a). In contrast to the cells with silenced c-Myc, removal of AHT efficiently restored inclusion formation (Fig. 1a), suggesting that suppression of c-Myc induces a persistence state in *Chlamydia*. Removal of AHT after 72 h still restored chlamydial inclusion formation, however, the HeLa cells started to lyse due to the long cultivation time (not shown). A physiological mechanism to induce chlamydial persistence is the exposure of infected cells to IFN-γ which causes the depletion of Trp via the induction of IDO^24^. Interestingly, IFN-γ-treated cells failed to stabilize c-Myc upon *Chlamydia* infection (Fig. 1b). In contrast, induction of persistence by antibiotic treatment, which targets the bacterial rather than host cell metabolism, did not alter c-Myc levels (Fig. 1b). Nevertheless, both modes of persistence induction caused a failure of the bacteria to produce infectious progenies (Extended data Fig. 1a).

**Figure 1:**
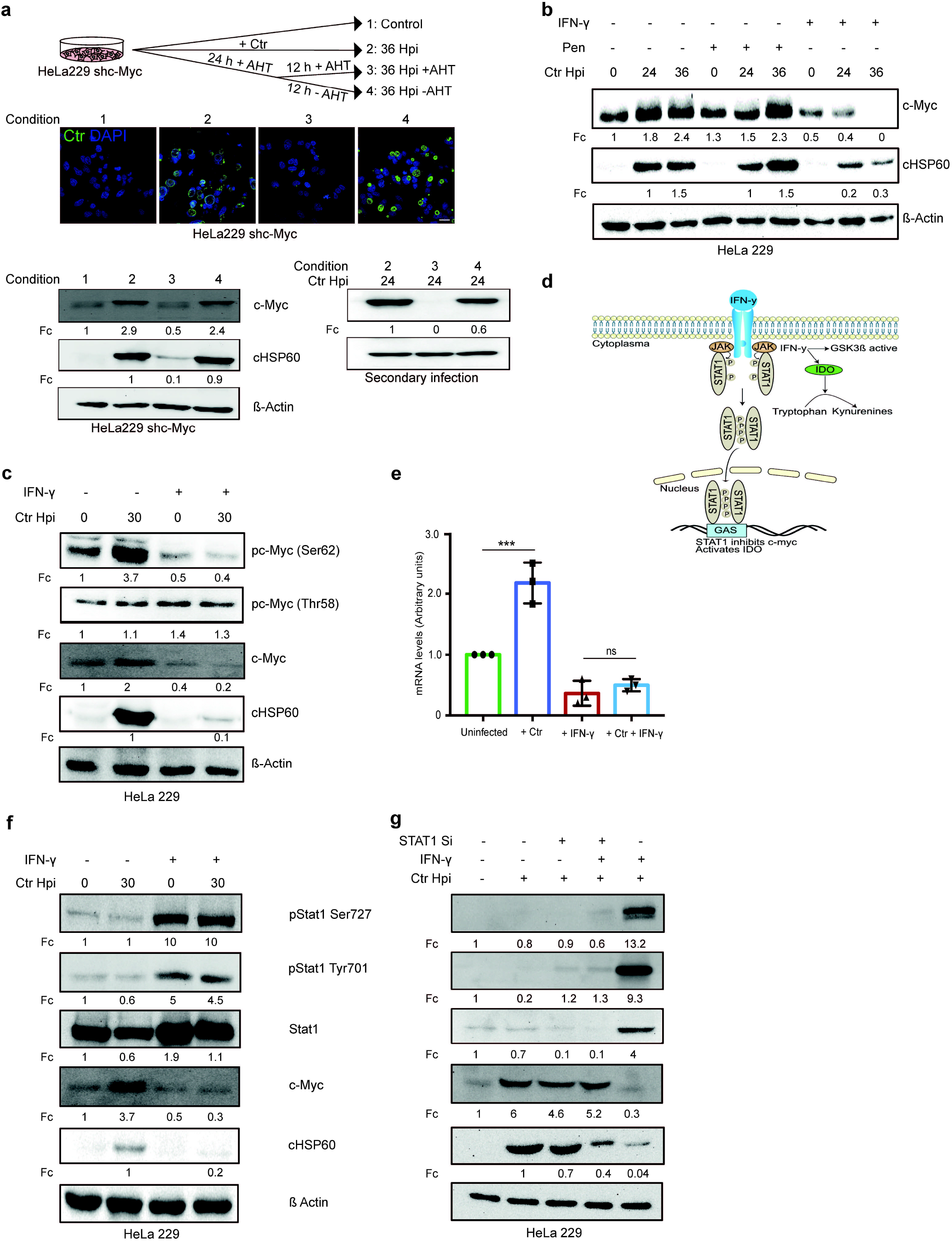
IFN-γ induces depletion of c-Myc and impairs chlamydial growth. **a.** HeLa 229 cells with an AHT-inducible expression of shc-Myc were infected with *Chlamydia* at MOI 1. The infected cells were either left untreated or treated with 1 ng/mL AHT to deplete c-Myc 8 h before infection. After 24 h of infection, AHT was removed to release c-Myc expression and the restoration of inclusion formation was tested. Cells were either fixed with 4% PFA after 36 hpi and immunostained for *Ctr* (cHSP60: green) and DNA (DAPI: blue) or lysed and analysed by Western blot in order to analyse the rescue. Additionally, infected cells were lysed to infect freshly plated HeLa 229 shc-Myc cells at 24 hpi and analysed via Western blot to investigate the formation of infectious progenies. The panel shows representative images (n=3). cHSP60 indicates *Chlamydia* infection and ß-Actin serve as the loading control (n=3). **b.** HeLa 229 cells were either left untreated or were pretreated for 2 h with 10 ng/mL IFN-γ or 1 unit of penicillin, infected with *Ctr* at MOI 1 for different time intervals and lysed for Western blot analysis. Bacterial load (cHSP60) and c-Myc levels were determined and ß-Actin served as loading control (n=3). **c.** HeLa 229 cells were either left untreated or were pre-treated for 2 h with 10 ng/mL of IFN-γ and infected with *Chlamydia (Ctr)* at MOI 1 and lysed at 30 h to perform Western blot analysis. Phosphorylated form of c-Myc at serine 62 (pc-Myc (Ser62)) and threonine 58 (pc-Myc (Thr58)), c-Myc and *Chlamydia* (cHSP60) were detected and quantified (Fc) (n=3). **d.** Cartoon depicting IFN-γ signalling. IFN-γ binds to the IFN-γ receptor that results in the phosphorylation of STAT1. pSTAT1 binds to the GAS sequence and blocks c-Myc transcription. IFN-γ can also induce indoleamine-2,3-dioxygenase (IDO) and thereby the degradation of L-tryptophan. **e.** HeLa 229 cells were either left untreated or treated with 10 ng/mL of IFN-γ and infected with *Ctr*. The cells were lysed and relative mRNA levels of c-Myc were determined by qPCR. GAPDH was used for normalization (n=3). *** indicates p value < 0.001 and ns indicates nonsignificant. **f.** HeLa 229 cells were either left untreated or were pre-treated for 2 h with 10 ng/mL of IFN-γ and infected with *Chlamydia (Ctr)* at MOI 1 and lysed at 30 hpi to study STAT1 signalling after *Ctr* infection (n=3). **g.** HeLa 229 cells were transfected with siRNA against STAT1 (+) or control (-) for 48 h and then infected with *Ctr* for 24 h. The cells were lysed and analysed by Western blot to investigate the STAT 1 signalling and infectivity of *Ctr* (n=3). For all blots shown in figure 1, *Chlamydia* load (cHSP60) and the respective host cell protein levels were quantified by normalization to ß-Actin and indicated as fold change (Fc).

c-Myc protein stability is regulated by two phosphorylation sites with opposing functions. Serine 62 phosphorylation (pS62) stabilizes c-Myc whereas threonine 58 phosphorylation (pT58) promotes c-Myc degradation^53^. IFN-γ treatment led to decreased phosphorylation of c-Myc at serine 62, a modification of c-Myc that prevents the ubiquitination and proteasomal degradation of the protein^54^, while phosphorylation at threonine 58 was unchanged (Fig. 1c). c-Myc was also depleted upon IFN-γ treatment of primary cells from human fimbriae (Fimb), although the concentration of IFN-γ required to achieve the same effect was five times higher than in HeLa 229 cells (10 ng/mL in HeLa 229 compared to 50 ng/mL in Fimb) (Extended data Fig. 1b, c). Similar results were obtained with a serovar D strain involved in STD (Extended data Fig. 1d).

IFN-γ is known to signal via the JAK-STAT pathway via phosphorylation of STAT1 and transcriptional regulation of gene expression (Fig. 1d). Treatment with IFN-γ led to the phosphorylation of STAT1 at serine 727 and tyrosine 701 (Fig. 1f) and prevented the induction of c-Myc protein (Fig. 1f) and mRNA expression (Fig. 1e) upon Chlamydia infection. To verify that interferon signals via STAT1 to transcriptionally deplete c-Myc, we used siRNA against STAT1 (Fig. 1g). STAT1 depletion rescued c-Myc levels and chlamydial growth in primary infections (Fig. 1g). When bacteria from IFN-γ treated and STAT1-depleted cells were transferred to fresh cells, infectious progeny could be recovered, indicating that STAT1 downregulates c-Myc and prevents chlamydial development (Extended data Fig. 1e).

The IFN-γ response in humans and mice is entirely different, limiting the use of murine systems as models for human pathogenic *C. trachomatis* infections^23^. We therefore established an IFN-γ-induced persistence model in human fallopian tube organoids. Healthy tissue obtained from patients that underwent hysterectomy was used to establish organoid cultures. These organoids obtained from five different patients were pre-treated with IFN-γ for 2 h and then infected with *Chlamydia* for 6 days (Fig. 2a, b). In this human infection model, IFN-γ treatment strongly reduced inclusion formation (primary infection: Fig. 2b) and production of infectious progeny (Fig. 2c, d; Extended Fig. 2). Moreover, the IFN-γ-STAT1 signalling axis was active in human organoids and efficiently prevented the induction of c-Myc upon infection (Fig. 2e), confirming the physiological significance of this system as a model of persistence infection.

**Figure 2:**
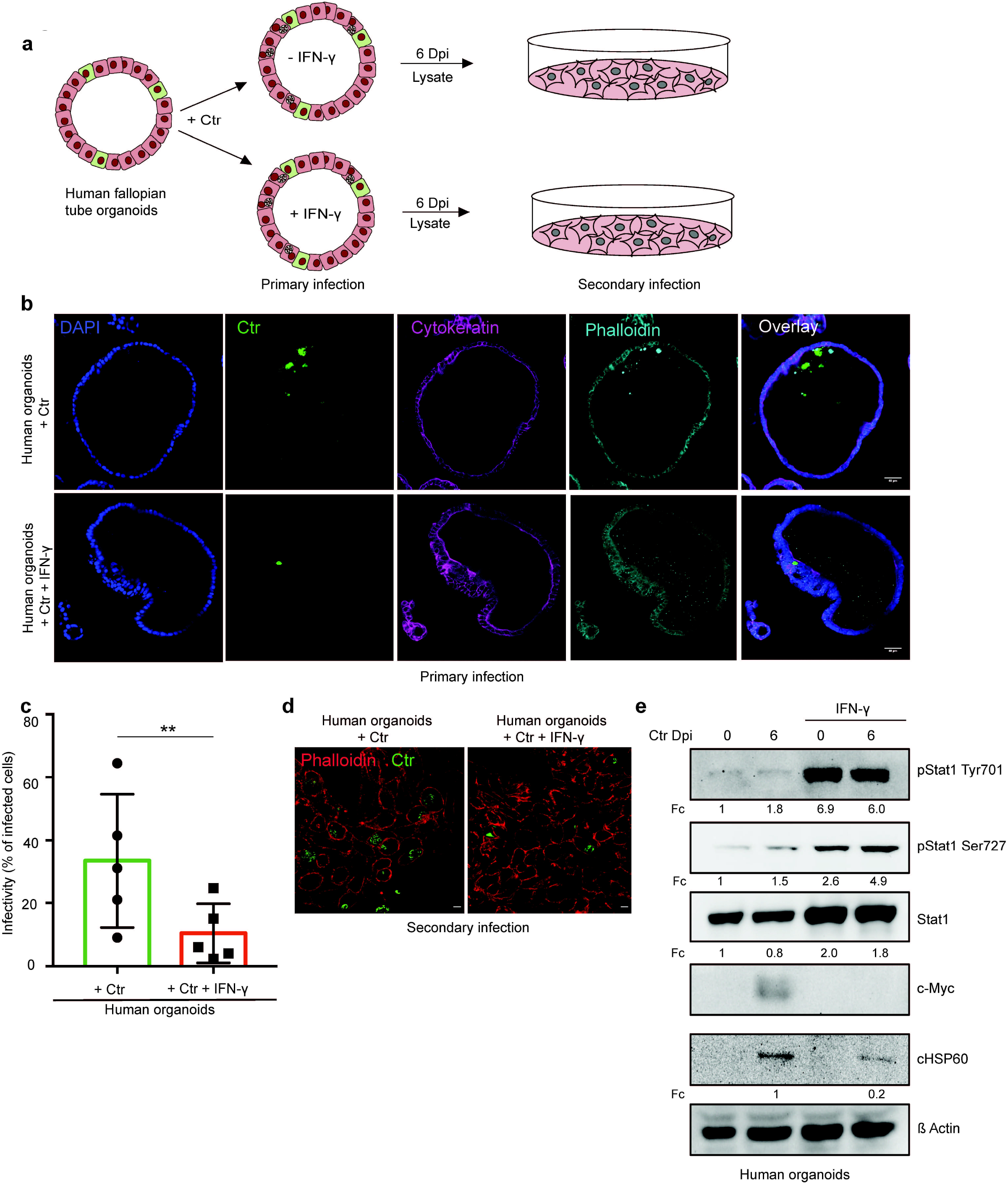
IFN-γ induces depletion of c-Myc and impairs chlamydial growth in human fallopian tube organoids. **a.** Cartoon depicting how the infectivity assay was performed in human fallopian tube organoids. Organoids (see Methods) were infected with *Ctr* and treated with or without IFN-γ for 6 days and lysed using glass beads. Dilutions of the supernatants were used to infect freshly plated HeLa 229 cells. **b.** Human fallopian tube organoids were infected with *Ctr* and treated with or without IFN-γ for 6 days. The organoids were fixed with 4% PFA and immunostained for DNA (DAPI: blue), *Ctr* (cHSP60: green), Cytokeratin (magenta), Phalloidin (light blue). The panel shows representative images from organoids derived from five patients. **c.** The infected organoids from (**b**) were lysed using glass beads, and dilutions of the supernatant were used to infect freshly plated HeLa 229 cells. The number of inclusions as shown in (**d**) were counted from 5 different patients and mean ± SD depicted in the graph (n=5). ** indicates p value < 0.01. **e.** Human fallopian tube organoids were infected with *Ctr* and treated with or without IFN-γ for 6 days. The organoids were lysed in 2x Laemmeli buffer and analysed by Western blotting. *Chlamydia* load (cHSP60) and STAT1 protein levels were quantified by normalization to ß-Actin and indicated as fold change (Fc).

### Expression of c-Myc rescues *Chlamydia* from IFN-γ-induced persistence

As we observed a key role of c-Myc in chlamydial persistence, we next asked if maintaining c-Myc levels can overcome IFN-γ-induced persistence. To address this question, we used a WII-U2OS cell line in which c-Myc expression is under the control of an AHT-inducible promoter. These cells were used as infection models for IFN-γ-induced persistence under constant c-Myc expression. Bacterial replication efficiency and the ability to produce infectious progenies were studied. Interestingly, constant expression of c-Myc efficiently suppressed IFN-γ-induced persistence (Fig. 3a) and supported the development of infectious progeny (Fig. 3b, c; Extended data Fig. 3a). We also tested the effect of c-Myc on persistence in organoids derived from human fallopian tubes. Organoids were transduced with lentivirus expressing c-Myc or GFP as control (Extended data Fig. 3b-d) and selected for puromycin resistance (Extended data Fig. 3e). These organoids were then infected with *Chlamydia* and treated with IFN-γ. Maintaining expression of c-Myc rescued the strong suppression of infectious progeny development by IFN-γ also in fallopian tube organoids (Fig. 3d, e), demonstrating that downregulation of c-Myc is essential for IFN-γ-mediated persistence in this human infection model.

**Figure 3:**
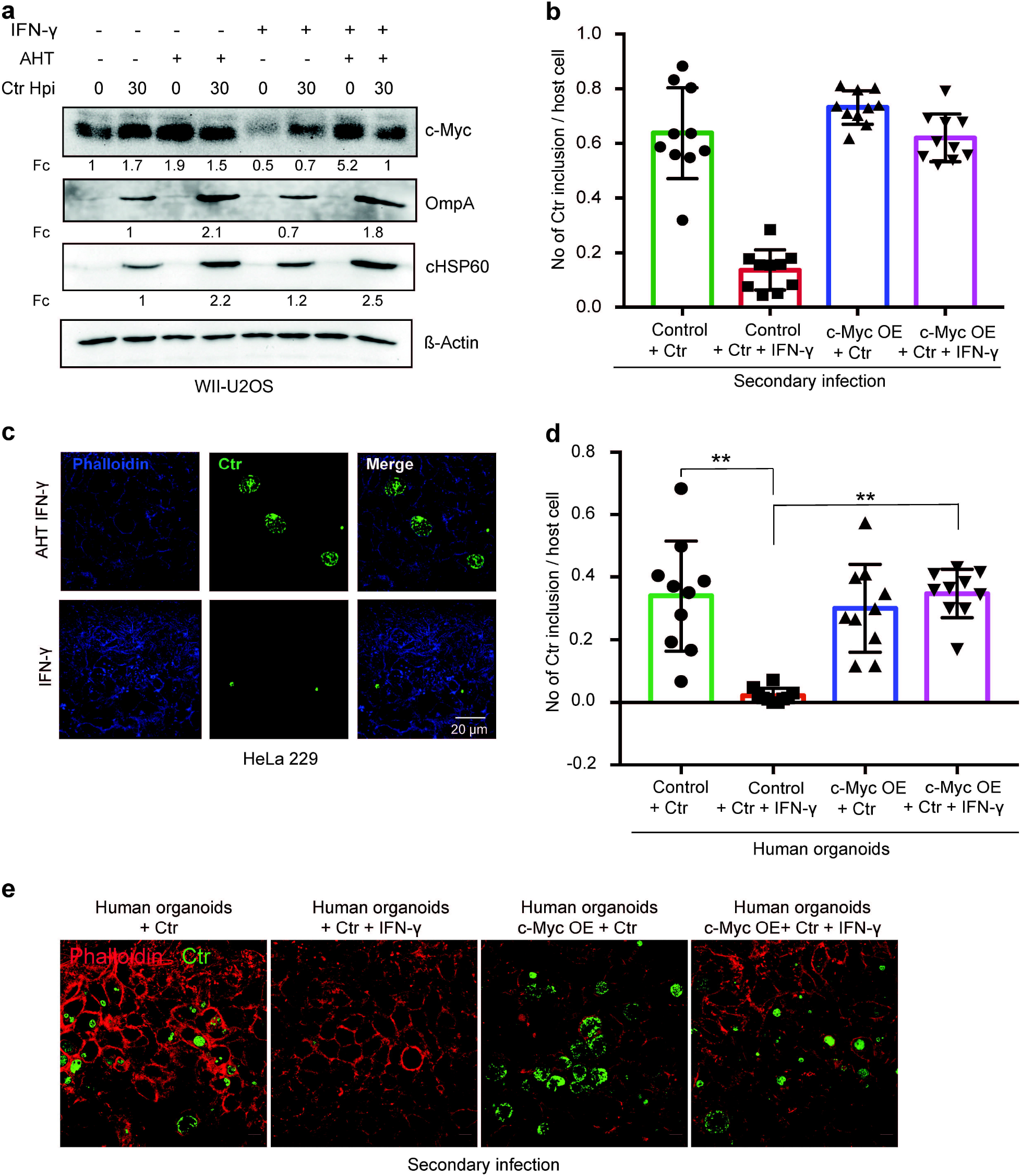
Expression of c-Myc rescues *Chlamydia* from persistence. **a.** WII-U2OS cells were induced with 100 ng/mL AHT for 2 h. Cells were left untreated or were pre-treated for 2 h with 10 ng/mL IFN-γ, infected with *Chlamydia (Ctr)* at MOI 1 and lysed at 30 hpi for Western blot analysis. **b.** The infected cells from (**a**) were lysed to infect freshly plated HeLa 229 cells. The numbers of inclusions were counted from the different conditions shown in (**a**). The mean ± SD are shown in the graph. *** p < 0.001, ns = not significant. **c.** For the same culture conditions as in (**a**), an infectivity assay was performed. The HeLa 229 cells were fixed with 4% PFA after 30 hpi and immunostained for *Ctr* (cHSP60: green) and actin (Phalloidin: blue) in order to analyse the rescue. **d.** From the experiment shown in (**e**) the infected/ IFN-γ treated organoids were lysed with glass beads and different dilutions of the supernatant was used to infect freshly plated HeLa 229 cells. The number of inclusions were counted from three different experiments and mean ± SD are shown in the graph. ** indicates a p value < 0.01. **e.** The organoids from (**Extended data Fig. 3b**) were infected with *Ctr* for 6 days with and without IFN-γ, were lysed using glass beads, and dilutions of the supernatant were used to infect freshly plated HeLa 229 cells for an infectivity assay. Cells were fixed with 4% PFA after 30 hpi and immunostained for *Ctr* (cHSP60: green) and actin (Phalloidin: red) in order to analyse the rescue.

### Both c-Myc and L-tryptophan are required to rescue *Chlamydia* from IFN-γ induced persistence

It has been shown previously that IFN-γ-mediated persistence can be overcome by the addition of exogenous Trp^18^. To investigate the importance of Trp in the context of IFN-γ-mediated regulation of c-Myc expression in our model system, HeLa 229 and human Fimb cells were treated with IFN-γ, infected with *Chlamydia* and provided with exogenous Trp. Subsequently, bacterial replication efficiency as well as production of infectious progenies was analysed by Western blotting. As expected, addition of Trp rescued chlamydial growth after IFN-γ treatment (Extended data Fig. 4a) and gave rise to infectious progenies (Fig. 4a; Extended data Fig. 4b). Surprisingly, addition of Trp also resulted in the stabilization of c-Myc, even without chlamydial infection (Extended data Fig. 4c).

**Figure 4:**
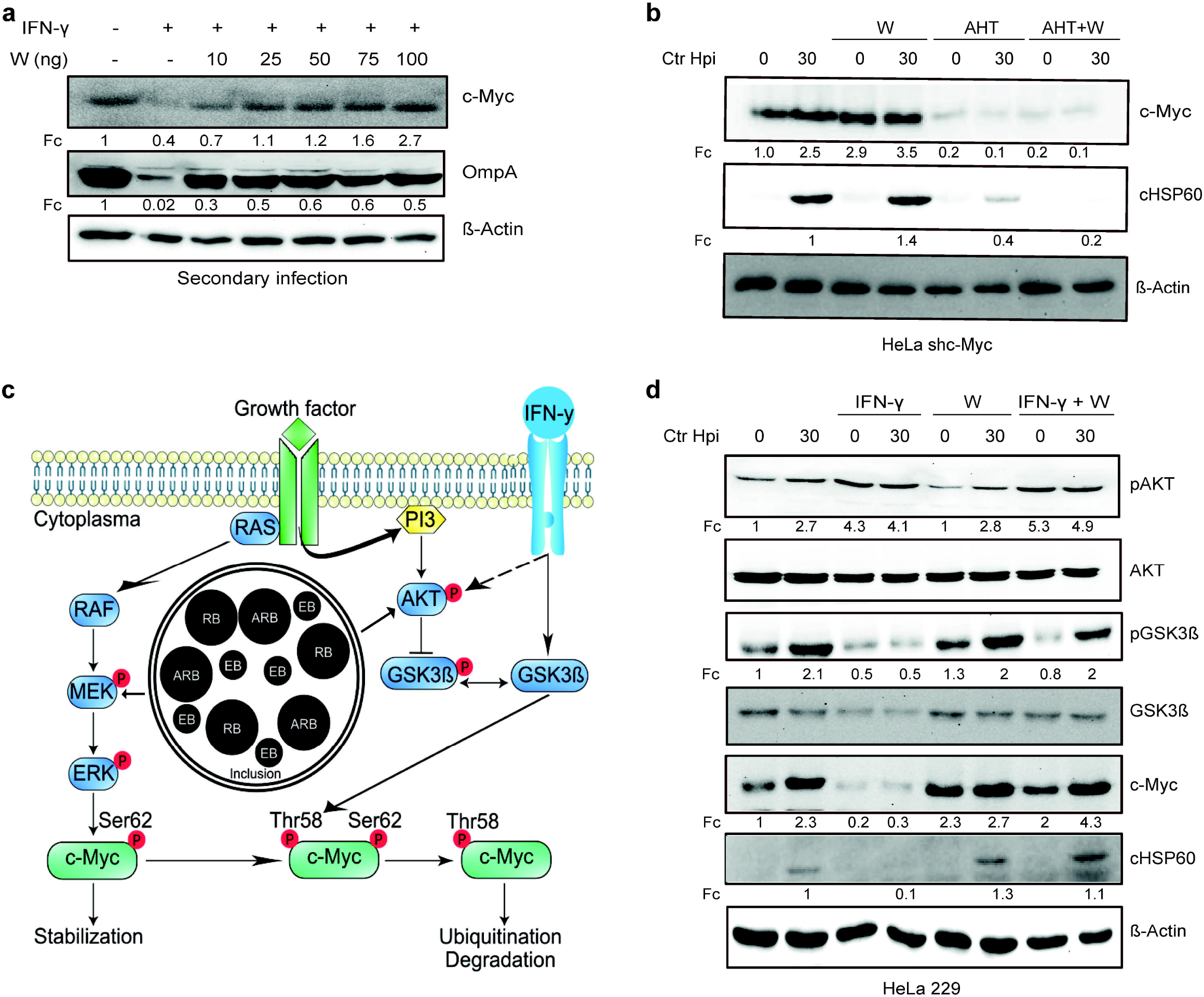
L-tryptophan activates the pGSK3ß-c-Myc axis and rescues chlamydial infection. **a**. HeLa 229 cells were treated with IFN-γ and different doses of L-tryptophan (W) (10-100 ng/mL). After 30 h the cells were lysed and used for Western blot analysis (n=3). **b.** HeLa 229 cells with an AHT-inducible expression of shc-Myc were either left uninfected or infected with *Chlamydia* at MOI 1 for 30 h. The infected cells were either left untreated or treated with L-tryptophan and 1 ng/mL AHT to deplete c-Myc. The cells were further analysed via Western blotting (n=3). **c.** Cartoon showing how IFN-γ signals lead to c-Myc depletion. Left side, active Phosphatidylinositol-3-kinase (PI3K) with inactive glycogen synthase kinase-3 (GSK3ß) leading to c-Myc stabilization. Right side, IFN-γ binding to its receptor, activates PI3K and serine-threonine protein kinase (AKT) and induces the dephosphorylation and activation of GSK3ß, leading to c-Myc depletion. *Chlamydia* infection activates the PI3K- and MEK/ERK-pathway. **d.** HeLa 229 cells were either left uninfected or infected with *Ctr* and treated with IFN-γ with or without L-tryptophan (W). The cells were analysed via Western blotting after 30 hpi (n=3). cHSP60 indicates *Chlamydia* infection and ß-Actin serve as the loading control.

All *C. trachomatis* strains are Trp auxotroph but the genital strains retain an L-tryptophan synthase (*trpB*), which uses exogenous indole provided by the microflora in the genital tract as a substrate to synthesize Trp^27,55,56^. Thus, we validated if exogenous indole recovers chlamydial growth and leads to infectious progenies. Intriguingly, indole addition resulted not only in suppression of persistence (Extended data Fig. 4d) and formation of infectious progeny (Extended data Fig. 4e), but also in the stabilization of c-Myc in both infected and non-infected cells (Extended data Fig. 4d).

Next, we tested whether Trp alone is sufficient to support *Chlamydia* development also in the absence of c-Myc. To address this question, we depleted c-Myc by AHT-inducible shRNA-mediated gene silencing and supplemented the medium with Trp as before. These cells were used for infection studies with *Chlamydia* and the formation of progeny was analysed via Western blot (Fig. 4b, Extended data Fig. 4f). Interestingly, the pathogen failed to develop in cells with reduced levels of c-Myc even in presence of excessive exogenous Trp (Fig. 4b). Furthermore, *Chlamydia* was also unable to establish a secondary infection under low c-Myc expression conditions, irrespective of Trp availability (Extended data Fig. 4f). Since excess Trp cannot overcome the suppression of c-Myc, we tested if the constant expression of c-Myc in a Trp free environment would be able to rescue chlamydial growth. Therefore, WII-U2OS cells were cultivated in Trp-free medium, induced with AHT, treated with IFN-γ and infected with *Chlamydia*. Bacterial replication efficiency and the ability to produce infectious progeny was examined by Western blotting. Neither chlamydial development nor a secondary infection could be observed (Extended data Fig. 4g, h). In addition, c-Myc was not stabilized upon chlamydial infection in the absence of Trp (Extended data Fig. 4g), suggesting that viable bacteria and Trp are required for the stabilization of c-Myc. These data support the role of c-Myc in Trp- or indole-mediated rescue from IFN-γ-induced chlamydial persistence.

### L-tryptophan addition leads to activation of pGSK3ß/c-Myc axis and restores chlamydial infection

Since we observed that Trp and c-Myc are both required for the development of *Chlamydia*, we next investigated the mechanism by which this amino acid regulates c-Myc level. IFN-γ, upon binding to its receptor, activates phosphatidylinositol-3-kinase (PI3K) and serine-threonine protein kinase (AKT) and induces the dephosphorylation and activation of glycogen synthase kinase-3 (GSK3ß)^57^. Dephosphorylated active GSK3ß leads to the phosphorylation of c-Myc at threonine 58, followed by its ubiquitination and proteasomal degradation^58^ (Fig. 4c). Both the MAPK and PI3K pathways activated during infection are critical for chlamydial development^59–62^. Therefore, the phosphorylation status of AKT and GSK3ß was examined in HeLa 229 and human Fimb cells, which were treated with IFN-γ and/or Trp and infected with *Chlamydia*. IFN-γ activated the PI3 kinase pathway as evident from the phosphorylation of AKT (Fig. 4d; Extended data Fig. 4i). Despite this increase in Akt phosphorylation, IFN-γ treatment strongly reduced the phosphorylation of GSK3ß and resulted in lower c-Myc levels (Fig. 4c, d; Extended data Fig. 4i). Surprisingly, the addition of Trp to HeLa or Fimb cells increased phosphorylation of GSK3ß, presumably resulting in its inactivation, and consequently led to an elevation of c-Myc levels (Fig. 4d; Extended data Fig. 4i). Taken together, these data demonstrate that Trp rescues chlamydial infection via the activation of the pGSK3ß-c-Myc axis.

### IFN-γ induced downregulation of c-Myc has a pleiotropic effect on host metabolism

Since c-Myc is centrally involved in regulating amino acid transport^63^, we asked if stabilized c-Myc increases also Trp uptake. In our previous RNA-seq analysis we observed the upregulation of the L-amino acid transporter Solute Carrier Family 7 Member 5 (LAT1/SLC7A5) in cells infected with *Chlamydia*^40^. LAT1 is a system L-amino acid transporter with high affinity for branched chain and bulky amino acids, including Trp^64^. Furthermore, the *LAT1* promoter has a binding site for c-Myc and it has been shown that overexpression of c-Myc leads to an increased expression of LAT1^64^. Therefore, the protein levels of LAT1 during a chlamydial infection and upon IFN-γ treatment were investigated by Western blotting. Accumulation of LAT1 protein was detected in a time-dependent manner in infected HeLa 229 cells (Fig. 5a). In contrast, LAT1 protein levels were strongly reduced in IFN-γ-treated cells, irrespective of infection (Fig. 5b). Following c-Myc expression, LAT1 levels in infected cells were rescued even in the presence of IFN-γ (Fig. 5c). Interestingly, the Trp-degrading enzyme IDO, which was strongly induced by IFN-γ treatment as expected, was only partially suppressed by c-Myc expression (Fig. 5c).

**Figure 5:**
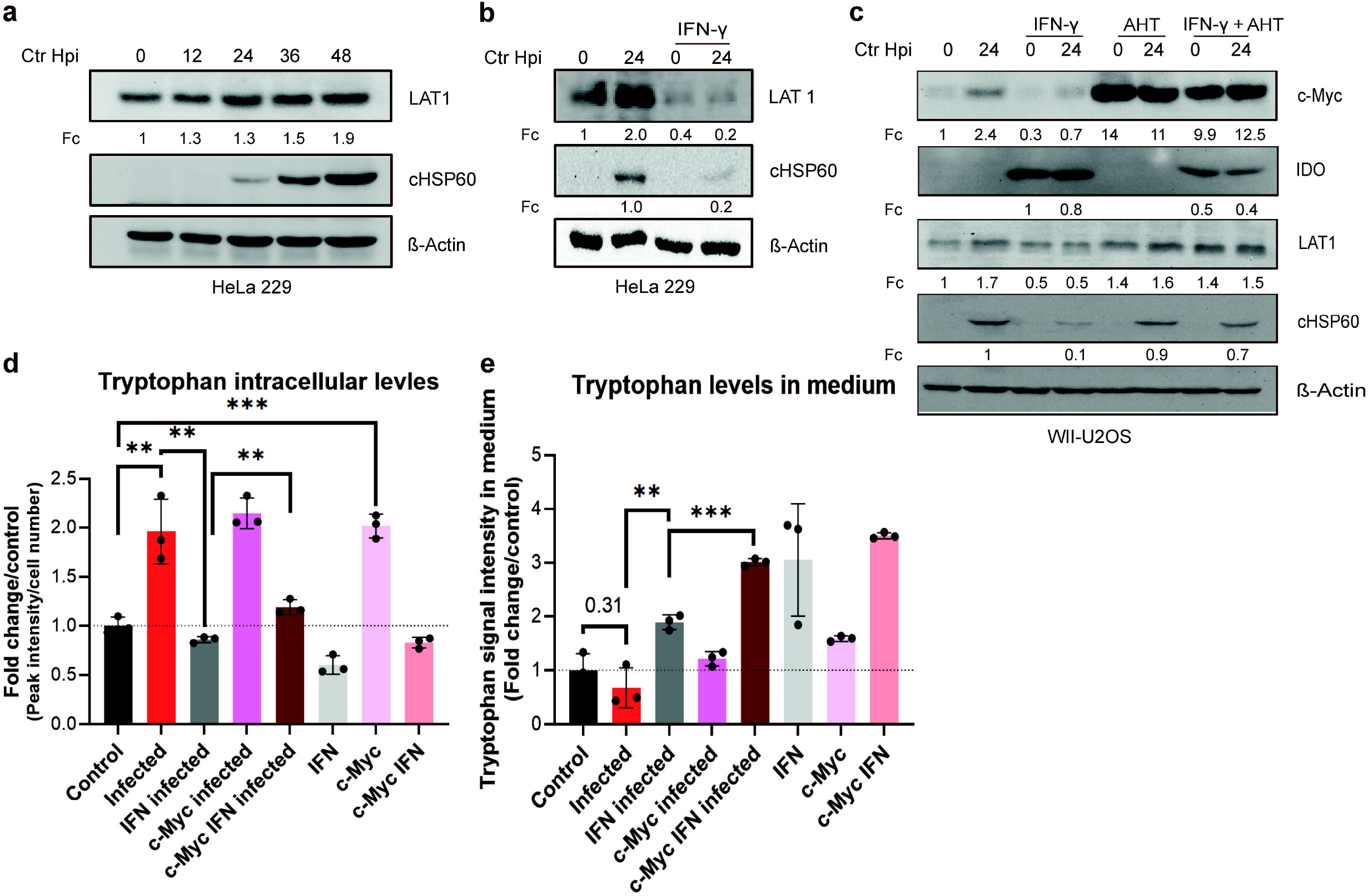
Influence of stabilized c-Myc on Trp uptake and metabolites. **a.** HeLa 229 cells were infected with *Chlamydia* (*Ctr*) for various time points. The cells were analysed via Western blotting for the levels of LAT1 (n=3). **b.** HeLa 229 cells were either left untreated or were pre-treated for 2 h with 10 ng/mL of IFN-γ, infected with *Ctr* at MOI 1 and lysed at 24 hpi to examine LAT1 regulation via Western blotting. **c.** WII-U2OS cells were induced with 100 ng/mL AHT for 2 h. Cells were left untreated or were pre-treated for 2 h with 10 ng/mL IFN-γ, infected with *Chlamydia (Ctr)* at MOI 1 and lysed at 24 hpi for Western blot analysis (n=3). **d.** WII-U2OS cells were either left uninfected or infected with *Ctr* at MOI 1 for 30 h. The infected cells were either left untreated or treated with just 10 ng/mL IFN-γ and 100 ng/mL AHT to induce expression of c-Myc. The cells were extracted, and metabolites were analysed by LC-MS. Data are presented as mean ± SD of triplicate wells. ** indicates a p value < 0.01, *** indicates a p value < 0.001. The intracellular levels of tryptophan are shown. **e.** WII-U2OS cells were either left uninfected or infected with *Ctr* at MOI 1 for 30 h. The infected cells were either left untreated or treated with just 10 ng/mL IFN-γ and 100 ng/mL AHT to induce over expression of c-Myc. The media in which the cells were grown was extracted and metabolites were analysed by LC-MS. Data are presented as mean ± SD of triplicate wells. ** indicates a p value < 0.01, *** indicates a p value < 0.001. The levels of tryptophan present in medium is shown.

To investigate how c-Myc restoration affects Trp metabolism in IFN-γ treated cells, we measured intracellular levels of Trp via LC-MS of WII-U2OS cells with an AHT-inducible c-Myc expression. This analysis showed that *Chlamydia* infection increases intracellular levels of Trp, while addition of IFN-γ results in decreased Trp levels (Fig. 5d). Interestingly, c-Myc expression in non-infected cells increased intracellular Trp to the levels of infected cells (Fig. 5d). However, in the presence of IFN-γ, expression of c-Myc only induced a small increase in intracellular Trp levels to about the same level observed in uninfected cells (Fig. 5d), indicating that c-Myc is not sufficient to prevent the effect of IFN-γ on Trp degradation. To also investigate whether induction of LAT1 by c-Myc affects intracellular Trp levels by increasing its uptake, we also measured Trp levels in the culture medium (Fig. 5e). Surprisingly, c-Myc expression had no major effect on Trp uptake in untreated and IFN-γ treated cells (Fig. 5e), suggesting that restoring Trp metabolism is not the main mechanism of overcoming chlamydial persistence downstream of IFN-γ signalling.

We next performed an unbiased metabolomics analysis. WII-U2OS cells were infected with *Chlamydia* and either induced with AHT, and/or treated with IFN-γ, and the resulting changes in metabolite levels were analysed by LC-MS. Quality controls and data normalization were performed and a principal component analysis (PCA) demonstrated the validity of the datasets (Extended data Fig. 5a). Interestingly, hierarchical clustering analysis revealed the grouping of all conditions permissive for chlamydial development (Fig. 6a), strongly suggesting that these metabolite profiles are indicative of productive chlamydial infection. Chlamydial development is favoured in an environment with high levels of amino acids (tyrosine, histidine, alanine, homoserine, methionine, lysine, phenylalanine, threonine, asparagine, glutamate), TCA (citrate, aconitate, malate, α-ketoglutarate) and urea cycle intermediates (ornithine, citrulline) (Fig. 6a).

**Figure 6:**
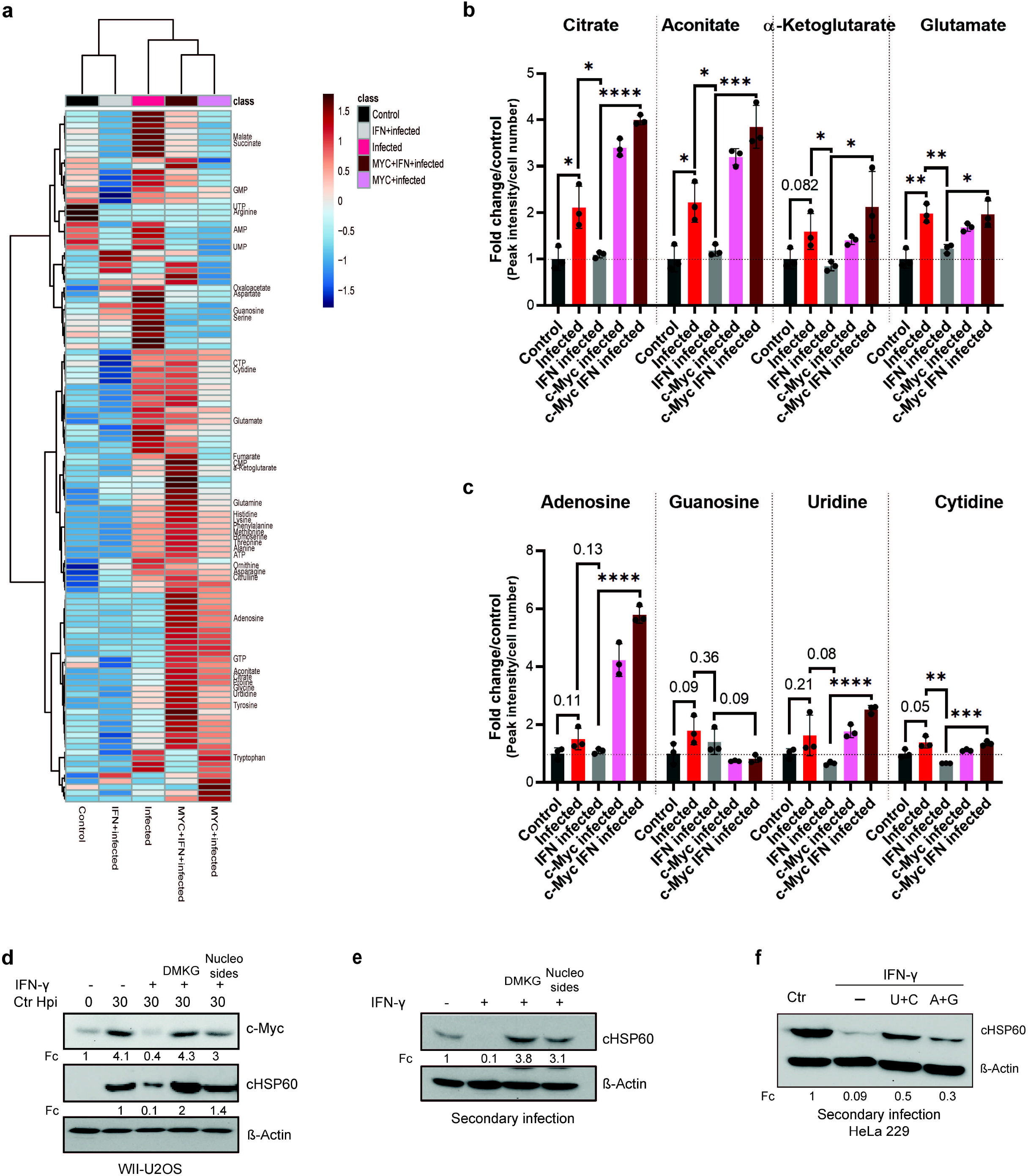
TCA intermediates and nucleosides can overcome IFN-γ-induced persistence. **a.** A Heatmap with hierarchical clustering of all metabolites detected by LC-MS analysis of (**Fig. 5d**). **b/c**. WII-U2OS cells were either left uninfected or infected with *Ctr* at MOI 1 for 30 h. The infected cells were either left untreated or treated with just 10 ng/mL IFN-γ and 100 ng/mL AHT to induce over expression of c-Myc. The cells were extracted, and metabolites were analysed by LC-MS. Data are presented as mean ± SD of triplicate wells. Intracellular levels of metabolites like citrate, aconitate, α-ketoglutarate, glutamate (**b**), nucleosides (adenosine, guanosine, cytidine and uridine) (**c**) were determined and quantified. * indicates a p value < 0.05, ** indicates a p value < 0.01, *** indicates a p value < 0.001, **** indicates a p value < 0.0001. **d.** Cells were left untreated or were pre-treated for 2 h with 10 ng/mL IFN-γ, 4 mM α-ketoglutarate (DMKG) or 100 μM nucleosides, infected with *Ctr* at MOI 1 and lysed at 30 hpi for Western blot analysis (n=3). The α-ketoglutarate was supplied as cell-permeable dimethyl ester. **e.** For the same culture conditions as in (**d**), an infectivity assay was performed. The cells were lysed at 48 hpi and dilutions of the resulting *Chlamydia* containing supernatant were added onto freshly plated WII-U2OS cells, which were then lysed at 30 hpi to examine infectious progenies via Western blotting (n=3). **f.** HeLa 229 cells were either infected with *Chlamydia* or infected and treated with IFN-γ and nucleosides (uridine/cytosine or adenosine/guanosine) were added. The cells were lysed at 48 hpi and dilutions of the resulting *Chlamydia* containing supernatant were added onto freshly plated HeLa 229 cells, which were then lysed at 30 hpi to examine infectious progenies via Western blot (n=3).

Pathway analysis revealed that nicotinate and nicotinamide metabolism is strongly regulated upon *Chlamydia* infection (Extended data Fig. 5b). Moreover, phenylalanine, tyrosine and tryptophan biosynthesis was among the significantly altered pathways with strongest impact upon infection, but also after IFN-γ treatment of infected and c-Myc overexpressing infected cells (Extended data Fig. 5b-d). Other modulated metabolic pathways included amino acid pathways that have been shown before to be regulated upon *Chlamydia* infection cells^46,65^. In addition, irrespective of IFN-γ treatment, arginine was one of the most depleted amino acids in infected cells (Extended data Fig. 6d).

### Alpha-ketoglutarate and nucleosides rescue *Chlamydia* from IFN-γ induced persistence

We next focused our attention on those metabolites that were induced by infection but reduced following IFN-γ treatment and restored by c-Myc overexpression as candidates that could be causally involved in *Chlamydia* persistence. *Chlamydia* infection resulted in a significant increase in several TCA cycle (related) intermediates, including aconitate, citrate, α-ketoglutarate and glutamate (Fig. 6b; Extended data Fig. 6a). Furthermore, the amino acids aspartate, serine and glycine, which function as important precursors for TCA cycle anaplerosis and nucleotide biosynthesis (Extended data Fig. 6b), were also significantly induced by *Chlamydia* infection, while glutamine showed a trend towards induction that failed to reach statistical significance. In contrast, levels of arginine were strongly reduced upon infection, while intracellular levels of the urea cycle metabolites ornithine and citrulline were significantly increased (Extended data Fig. 6d).

Remarkably, treatment of infected cells with IFN-γ lowered the induction of glutamate as well as aspartate, serine and glycine. Moreover, levels of most TCA cycle metabolites, in particular aconitate, citrate, α-ketoglutarate, succinate, fumarate and malate, were also reduced upon IFN-γ treatment, suggesting that reprogramming of host cell metabolism is a major part of the IFN-γ response (Fig. 6a, b). Interestingly, re-expression of c-Myc restored levels of the TCA cycle metabolites citrate, aconitate and α-ketoglutarate as well as the amino acids glutamine, glutamate and glycine in IFN-γ treated infected cells (Fig. 6a, b; Extended data Fig. 6b), suggesting that these metabolites are required for *Chlamydia* development.

We also investigated intracellular levels of purine and pyrimidine nucleotides and nucleosides as well as intermediates of nucleotide metabolism, as *Chlamydia* is an auxotroph for nucleotides. Interestingly, IFN-γ treatment significantly lowered the amounts of ATP and CTP as well as AMP, GMP, UMP and cytidine (Extended data Fig. 6e, f; Fig. 6c). Moreover, several other metabolites involved in nucleotide metabolism showed a trend towards reduced abundance in IFN-γ treated infected cells (Extended data Fig. 6e, f; Fig. 6c). This reduction in essential precursors for chlamydial DNA replication may explain why *Chlamydia* enter persistence in the presence of IFN-γ. In addition, overexpression of c-Myc enhanced levels of nucleoside triphosphates and strongly increased intracellular levels of adenosine, uridine and cytidine in IFN-γ treated cells (Fig. 6c; Extended data Fig. 6e).

Based on the observation that IFN-γ leads to a reduction in TCA cycle intermediates and nucleotides in host cells, we investigated whether chlamydial growth after IFN-γ induced persistence could be overcome by the supplementation of specific metabolic precursors. We therefore treated WII-U2OS and HeLa 229 cells with IFN-γ, followed by *Chlamydia* infection and addition of the cell-permeable dimethyl ester of α-ketoglutarate (DMKG) or a mixture of nucleosides (A, C, G, U). In both cell lines, the growth of *Chlamydia* was rescued by the addition of either DMKG or nucleosides and both conditions produced infectious progenies (Fig. 6d-f). Moreover, restoration of chlamydial development could also be achieved by the sole addition of pyrimidine nucleosides (U+C), and to a minor extend also purine nucleosides (A+G) (Fig. 6f). Taken together, these results indicate that IFN-γ broadly alters the metabolism of host cells to limit the availability of metabolic precursors for pyrimidine and purine biosynthesis to promote chlamydial persistence.

## Discussion

Persistent and recurrent infections are an important cause of excessive inflammation and tissue damage in the fallopian tube resulting in infertility and ectopic pregnancy^66,67^. Chlamydial infection of the epithelial cells lining the genital tract leads to the secretion of cytokines, like IL-8^68^ and GM-CSF^69^, which attract myeloid and lymphoid cells towards the site of infection. This local immune response leads to chronic inflammation and may promote the development of malignancy including ovarian cancer^70^. In addition, IFN-γ secreted by infiltrating T cells and NK cells provokes persistence of *Chlamydia* in epithelial cells. The current model of the immune defence elicited by IFN-γ in human cells centres around the induction of IDO and the consecutive degradation of Trp^17,18^. *Chlamydia* is auxotroph for Trp and enters persistence if this amino acid is degraded in the infected cell.

Our detailed molecular analysis of the IFN-γ induced metabolic alterations in the host cells challenges this model and shifts the focus to the key transcription factor and proto-oncogene c-Myc as a central regulator of *Chlamydia* persistence. We demonstrate that IFN-γ acts via the GSK3ß-STAT1 axis to deplete c-Myc levels and that constitutive expression of c-Myc is sufficient to rescue *Chlamydia* from persistence induced by IFN-γ (Fig. 3). Interestingly, this effect was also reproduced in human fallopian tube organoids, a newly developed physiologically relevant model for *Chlamydia* infection.

The pathway governing persistence uncovered in our study was indeed dependent on the levels of Trp (Fig. 4), since *Chlamydia* failed to grow in the absence of this amino acid even in the presence of continuous c-Myc expression (Fig. 4; Extended date Fig. 4). Conversely, the expression of c-Myc alone without Trp provision was not sufficient to achieve normal chlamydial growth (Fig. 4; Extended data Fig. 4). Nevertheless, our results clearly show that both Trp and c-Myc are required for efficient bacterial replication. *Chlamydia* utilizes Trp to synthesize proteins, like the outer membrane protein MOMP, a Trp rich polypeptide. Our findings that Trp but also indole, which *C. trachomatis* can use as a substrate for Trp synthesis^27,55,56^, affects the level of c-Myc was unexpected (Extended data Fig. 4). It was shown previously that glutamine deprivation lowers levels of adenosine nucleotides and suppresses c-Myc in cancer cells by a mechanism dependent on the c-Myc 3’UTR^71^. We show here that Trp induces the phosphorylation and inactivation of GSK3ß and thereby prevents the degradation of c-Myc by the ubiquitin proteasome system (Fig. 4; Extended data Fig. 4). Phosphorylation-dependent inactivation of GSK3ß also occurs during normal chlamydial infection and causes the accumulation of c-Myc protein, as GSK3ß leads to a destabilization of c-Myc by phosphorylation on the threonine 58 and proteasomal degradation^58,72^. This finding suggests that Trp activates the pGSK3ß/c-Myc axis in order to rescue *C. trachomatis* from persistence. Interestingly, c-Myc transcriptionally activates expression of the tryptophan transporter LAT-1, thereby increasing the uptake of Trp as part of a positive feed-back regulation. Moreover, Trp depletion induced by IFN-γ signalling via STAT1 leads to loss of c-Myc expression, indicating that this amino acid functions as a central regulator of host cell metabolism and determines the decision between chlamydial development vs. the induction of persistence.

Detailed analysis of supernatants and extracts of infected cells also pointed to a broader impact of IFN-γ treatment on host cell metabolism. While Trp levels were significantly increased in infected cells and strongly reduced by IFN-γ treatment, constitutive expression of c-Myc failed to restore Trp levels in IFN-γ treated cells despite preventing *Chlamydia* persistence (Fig. 5d, e). This already indicated that additional metabolic pathways regulated by c-Myc expression may be responsible for preventing persistence induced by IFN-γ. Unbiased metabolomics analysis of IFN-γ treated, infected and c-Myc-expressing cells and subsequent hierarchical clustering analyses revealed a grouping of all cells permissive for chlamydial replication (Fig. 6a; infected, c-Myc infected, c-Myc IFN-γ infected), suggesting that this is the metabolic profile required for permissive chlamydial replication.

Since c-Myc acts as a major metabolic regulator^73,74^, we systematically investigated the c-Myc-dependent alterations in metabolite levels in response to bacterial infection and IFN-γ treatment. This analysis revealed that several intermediates of the TCA cycle, including citrate, aconitate and α-ketoglutarate, as well as glutamate, were increased upon chlamydial infection, but significantly depleted upon IFN-γ-treatment and restored when c-Myc was reexpressed (Fig. 6). In addition, several nucleotides, in particular ATP and CTP were among the top metabolites upregulated upon chlamydial infection. Moreover, glutamate and alanine, all upregulated by infection, repressed in response to IFN-γ treatment and restored by c-Myc expression, are central metabolites for *Chlamydia*, since they serve as precursors for cell wall biosynthesis^75^. Interestingly, we could observe a strong increase in the levels of citrate in c-Myc overexpressing cells (Fig. 6b). Citrate is required for the synthesis of fatty acids, which are scavenged by the bacteria from the host cell, as *Chlamydia* lack citrate synthase, aconitase and isocitrate dehydrogenase and thus have only an incomplete TCA cycle^76,77^. The finding of others that reduced uptake of glucose may play a role in IFN-γ induced chlamydial persistence supports our model, since glycolysis fuels the TCA and nucleotide biosynthesis^78^.

*Chlamydia*-infected cells showed a drastic reduction in the levels of arginine (Extended data Fig. 6d). However, arginine levels were not restored upon c-Myc re-expression in IFN-γ treated infected cells. It is possible that arginine is shuttled into the urea cycle, as we observed significantly higher levels of ornithine and citrulline (Extended data Fig. 6d), which can be used for polyamine biosynthesis, that stabilize DNA and are essential for cell proliferation^79^. Further work is required to elucidate the exact role of arginine metabolism in chlamydial infection.

Here we show a central role of c-Myc in the control of the host cell metabolism during IFN-γ-mediated innate immune defence against *C. trachomatis* infection. The effect of IFN-γ-mediated c-Myc downregulation included substantial remodelling of host cell metabolism, including reduced abundance of TCA cycle intermediates, which are usually replenished via c-Myc-dependent glutaminolysis and anaplerosis. Thus, the enhanced production of amino acids, nucleotides and lipids, and, additionally, TCA cycle intermediates transferred from the host cell to the bacteria^46,80^ may all be affected by IFN-γ treatment^38^. The central role of host TCA cycle and nucleotide biosynthesis for chlamydial development is also supported by our finding, that supplementing cell-permeable α-ketoglutarate or nucleosides can overcome the IFN-γ-induced persistent state of *Chlamydia* (Fig. 6). It can therefore be concluded that c-Myc is required for *C. trachomatis* to induce metabolic reprogramming of the host cell to avoid nutrient shortage during replication and prevent the induction of persistence. IFN-γ depletes the essential amino acid Trp and causes a reduction in the levels of c-Myc, which affects the metabolic state of the host cell in a way that prevents chlamydial replication and induces persistence. It is very likely, that similar mechanisms are also involved in IFN-γ induced persistence of other intracellular bacteria^81^. In addition, IFN-γ also restricts the infection of several viruses^82,83^. Since c-Myc activation and metabolic reprogramming of cells is a prerequisite for the replication of several viruses^84,85^, our findings may be generally relevant for the IFN-γ dependent innate immune defence against infection.

## Methods

### Cell lines and bacteria

Human Fimb cells (epithelial cells isolated from the fimbriae of patients undergoing hysterectomy), HeLa 229 cells (ATCC® CCL-2.1™) and HeLa 229 pInducer11 shc-Myc cells were cultured in 10% (v/v) heat inactivated FBS (Sigma-Aldrich) RPMI1640 + GlutaMAX™ medium (Gibco™). WII - U2OS cells^86^ were maintained in high-glucose DMEM (Sigma-Aldrich) with 10% (v/v) heat inactivated FBS. All cell lines were grown in a humidified atmosphere containing 5% (v/v) CO_2_ at 37°C. In this study *Chlamydia trachomatis* serovar L2/434/Bu (ATCC® VR-902B^™^) and *Chlamydia trachomatis* serovar D/UW-3/Cx (ATCC® VR-885^™^) were used, and cultured and purified as published previously^87^. In brief, *Chlamydia* were propagated in HeLa 229 cells at a multiplicity of infection (MOI) of 1 for 48 h. Cells were mechanically detached and lysed using glass beads (3 mm, Roth). Low centrifugation supernatant (10 min at 2,000 g at 4°C) was transferred to high speed centrifugation (30 min at 30,000 g at 4°C) to pellet the bacteria. The pellet was washed and resuspended in 1x SPG buffer (7.5% sucrose, 0.052% KH_2_PO_4_, 0.122% Na_2_HPO_4_, 0.072% L-glutamate). Aliquots were made, stored at −80°C and the bacteria were titrated to have MOI of 1 to be used further in the experiments. Infected cells were incubated in a humidified atmosphere with 5% (v/v) CO_2_ at 35°C. The cell lines as well as the *Chlamydia* used in this study were tested to be free of *Mycoplasma* via PCR.

### Infectivity assay

For primary infection, control or IFN-γ-treated cells were infected with *Chlamydia trachomatis* at MOI 1 for 30-48 h. The cells were lysed with glass beads and freshly plated cells were infected with different dilutions (referred to as secondary infection). Lysate of the secondary infection were taken to determine the infectivity via Western blotting against chlamydial HSP60.

### Induction and chemicals

In order to induce c-Myc over expression in WII-U2OS cells or c-Myc knock down HeLa 229 pInducer11 shc-Myc cells, medium was changed to 10% (v/v) heat inactivated FBS DMEM or RPMI accordingly and 100 ng/mL AHT (Anhydrotetracycline hydrochloride; Acros) was added. After 2 h of induction, cells were infected with *Chlamydia trachomatis* and incubated for further 24 to 36 h. Interferon-gamma (Gibco™, Merck Millipore), L-tryptophan (Sigma-Aldrich) and Indole (Sigma-Aldrich) were added to cells as needed 2 h before infection.

### Western blot and antibodies

For Western blot analysis, cells were directly lysed with 2x Laemmli (10% 1.5M Tris-HCl pH 6.8, 4% SDS, 30% glycerol and 1.5% ß-mercaptoethanol) buffer on ice. Protein samples were separated with a 10% SDS-PAGE (Peqlab) and transferred onto a PVDF membrane (Sigma-Aldrich) via a semi-dry blotter (Peqlab) (2 hours at 1 mA per cm^2^). The membrane was blocked in 5% (w/v) non-fat dried milk powder in 1x Tris-buffered saline with 0.5% Tween20 (Sigma-Aldrich) for 1 h and then incubated in the appropriate primary antibody over night at 4°C; c-Myc, anti c-Myc phospho-T58 and anti c-Myc phospho-S62 were purchased from Abcam. Phospho-Akt (Ser473), Akt (pan), phospho-Erk, Erk, phospho-GSK-3ß (Ser9), GSK-3ß (3D10), phospho-Stat1 (Ser727), phospho-Stat1 (Tyr701) and Stat1 were obtained from Cell Signaling. ß-Actin was purchased from Sigma-Aldrich. Chlamydial HSP60 was obtained from Santa Cruz and OmpA was self-made. Proteins were detected with corresponding horseradish peroxidase (HRP)-conjugated secondary antibody (Santa Cruz), using homemade ECL solutions and Intas Chemiluminescence Imager.

### Immunofluorescence analysis

The cells were seeded on cover slips and infected with *Chlamydia trachomatis* serovar L_2_ at MOI 1 for indicated time points. Before fixation with 4% PFA/Sucrose (Roth), the cells were washed with DPBS (Gibco™). Fixed cells were permeabilized with 0.2% Triton-X-100 (Sigma-Aldrich) in 1x DPBS for 30 min, blocked with 2% FBS in 1x DPBS for 45 min and incubated with primary antibodies for 1 h at room temperature. Primary antibodies were diluted in 2% FBS in 1x DPBS; chlamydial HSP60 (Santa Cruz, 1:500) and Phalloidin (Thermo Fischer). Samples were washed and incubated with fluorescence dye conjugated secondary antibodies (Dianova) for 1 h in the dark at room temperature. Cover slips were mounted onto microscopy slides using mowiol, slides were air dried for at least 24 h and examined using a LEICA DM2500 fluorescence microscope. The images were analysed with LAS AF program and Image J software.

### Metabolic profiling

For this study, 10^6^ WII-U2OS cells per well were seeded in triplicates, either uninfected or infected with *Chlamydia trachomatis* serovar L_2_ for 30 h, induced and treated with IFN-γ. After the respective time, medium was collected, snap frozen in liquid nitrogen, and the cells were washed with ice cold 154mM ammonium acetate (Sigma) and snap frozen in liquid nitrogen. The cells were harvested after adding 480ul cold MeOH/H_2_O (80/20, v/v) (Merck) to each sample containing Lamivudine (Sigma) standard (10μM). The cell suspension was collected by centrifugation and transferred to an activated (by elution of 1mL CH_3_CN (Merck)) and equilibrated (by elution of 1mL MeOH/H_2_O (80/20, v/v)) C18-E SPE-column (Phenomenex).

The eluate was collected and evaporated in SpeedVac and was dissolved in 50 μL CH_3_CN/5mM NH_4_OAc (25/75). Each sample was diluted 1:1 (cells) or 1:5 (medium) in CH_3_CN. 5 μL of sample was applied to HILIC column (Acclaim Mixed-Mode HILIC-1, 3μm, 2.1 * 150 mm). Metabolites were separated at 30°C by LC using a DIONEX Ultimate 3000 UPLC system (Solvent A: 5mM NH4OAc in CH_3_CN/H2O (5/95), Solvent B: 5mM NH4OAc in CH_3_CN/H2O (95/5); Gradient: linear from 100% B to 50% B in 6 min, followed by 15 min const. 40% B, then returning to 100% B within 1 min) at a flow rate of 350 μL/min. After chromatographic separation, masses (m/z) were acquired using a Q-Exactive instrument (Thermo Fisher Scientific) in positive and negative ionization mode in the scan range between 69-1000 m/z with a resolution of 70,000. AGC target was set to 3×10^6 and maximum injection time was set to 200 ms. Sheath, auxiliary and sweep gas were set to 30, 10 and 3, respectively, and spray voltage was fixed to 3.6 kV. S-lens RF level was set to 55.0, the capillary and the Aux gas heater were heated to 320°C and 120°C, respectively. Peak determination and semi-quantitation were performed using TraceFinder™ Software. Obtained signal intensities were normalized to the internal standard (lamivudine) and a cell number, which was determined by crystal violet staining. In brief, for the crystal violet staining cells were fixed with 4% PFA/Sucrose (Roth), stained with 0.1% crystal violet (Merck) dissolved in 20% ethanol (Roth), washed with water, dried overnight and the absorbance was measured at 550nm. The pellet of the cell samples was dried, resuspended in 0.2M sodium hydroxide (Roth), cooked for 20 min at 95°C and absorbance was measured at 550nm. Statistical analysis was performed using Prism GraphPad. Hierarchical clustering, principal component analyses (PCA) and pathway analyses were done and resulting plots were generated by MetaboAnalyst 4.0^88^.

### Generation of human fallopian tube organoids

Generation of human organoids was adapted from^89^. Fallopian tube tissue was obtained from patients who underwent hysterectomy. The tissue was prepared and processed within 2 h. Briefly, tissue samples were washed with DPBS (Gibco) and placed into a sterile Petri dish (Corning) where they were cut into small pieces. Then, on the top of the minced tissue, a glass slide (VWR) was placed and strongly pressed down to obtain smaller pieces. The cells were washed with DPBS, placed into a 15 mL falcon tube and centrifuged at 1,000 g for 10 min. The supernatant was removed, the pellet was resuspended in Matrigel (Corning) and plated in 50 μL drop in wells in a 24-well plate. The plate was carefully transferred to 37°C incubator to allow the Matrigel to get solidified for 20 min following the addition 500 μL/well of pre-warmed media (DMEM advanced (Sigma), Wnt (25%), R-Spondin (25%), Noggin (10%), B27 (2%; Thermo Scientific), Nicotinamide (1 mM; Sigma), human EGF (50 ng/mL; Thermo Scientific), FGF (100 ng/mL; Thermo Scientific), TGF-ß inhibitor (0.5 mM; Tocris), Rock inhibitor (10 mM; Abmole Bioscience).

### Splitting organoids

Approximately in 7 days, the Matrigel drop was carefully resuspended in cold DMEM medium and centrifuged at 1,000 g at 4°C for 5 min. The supernatant was discarded and 50 μL Matrigel was added and further processed as explained above.

### Infectivity assay in organoids

For infection with *C. trachomatis* mature organoids were released from a confluent Matrigel drop by resuspending it with ice-cold DPBS (Gibco). The suspension was collected in a low-binding Eppendorf tube and infected with *C. trachomatis* L_2_ (5 x 10^5^ IFU). The suspension was mixed and placed on ice for 30 min following centrifugation. 50 μL of Matrigel was added to each tube and seeded into a 24-well plate (Corning) with following 20 min incubation at 37°C to allow the Matrigel drop solidify. Six days post infection the organoids were fixed with 4% PFA and used for immunostaining. In addition, infected organoids were lysed with glass beads and different dilutions were used to infect freshly plated HeLa 229 cells to analyse the infectivity of the progeny.

## Acknowledgement

We thank Dr. Jörg Wischhusen for providing the human Fimb cells and Dr. Francesca Dejure for suppling us with WII-U2OS cells. We thank Dr. Andreas Demuth for critically reading the manuscript. This work was supported by the Deutsche Forschungsgemeinschaft (DFG) priority program GRK2157 “3D Tissue Models for Studying Microbial Infections by Human Pathogens” and the European Research Council (grant no. ERC-2018-ADG/NCI-CAD) to T.R.

## Author contribution

N.V., K.R. and T.R. conceived and designed the study; N.V., K.R., N.K. and S.J.R. performed the experiments; S.J.R, N.V, L.S., W.S. and A.S. performed the mass spec analysis. N.V., T.R and K.R. wrote the manuscript, further edited by S.J.R., L.S., and A.S.; T.R, financially supported the project. K.R. supervised the work.

---

**Extended data Figure 1: IFN-γ induces c-Myc depletion and impairs chlamydial growth.**

**a.** An infectivity assay was performed in HeLa 229 cells to test for infectious progenies after addition of IFN-γ (10 ng/mL) or penicillin (1 unit). Lysate were subjected to Western blot analysis and bacterial load (cHSP60) was quantified by normalizing to ß-Actin levels. **b/c.** HeLa 229 cells/ human Fimb cells were treated with different doses of IFN-γ (10-50 ng/mL) and either left uninfected or infected with *Ctr* for 30/24 h. The cells were lysed and analysed via Western blotting (n=3). **d.** HeLa 229 cells were treated with 10 or 50 ng/mL of IFN-γ and infected with *Ctr* D serovar. After 48 h the cells were lysed with glass beads and the supernatant was used to infect freshly plated HeLa 229 cells. After 30 h the cells were lysed and analysed via Western blotting. Bacterial load was determined by quantifying cHSP60 levels and ß-Actin serves as loading control. **e.** Infectivity assay of the experiment shown in (**Fig. 1g**). The cells were lysed at 48 hpi and dilutions of the resulting *Chlamydia-containing* supernatants were added onto fresh HeLa 229 cells, which were then lysed at 30 hpi for Western blot analysis (n=3).

**Extended data Figure 2: IFN-γ-induced persistence of *Chlamydia trachomatis* in human organoids.**

Human fallopian tube organoids were infected with *Ctr* and treated with or without IFN-γ for 6 days. The infected organoids were lysed using glass beads, and dilutions of the supernatant were used to infect freshly plated HeLa 229 cells for an infectivity assay. Cells were fixed with 4% PFA after 30 hpi and immunostained for *Ctr* (cHSP60: green) and actin (Phalloidin: red). The data from 5 different patients are presented here.

**Extended data Figure 3: Expression of c-Myc rescues *Chlamydia* from persistence.**

**a.** For the same culture conditions as in (**Fig. 3a**), an infectivity assay was performed. The cells were lysed at 48 hpi and dilutions of the resulting *Chlamydia-containing* supernatant were added onto fresh HeLa 229 cells, which were then lysed at 30 hpi to examine infectious progenies. The cHSP60 detect *Chlamydia* infection and the ß-Actin serves as loading control (n=3). **b.** Human fallopian tube organoids were transduced with either lentivirus expressing c-Myc or GFP as control (see Methods). The puromycin selected organoids where lysed and analysed via Western blotting. ß-Actin serves as loading control. **c/d.** HeLa 229 cells were transduced with lentivirus expressing c-Myc (**c**) or GFP (**d**). The cells were analysed by Western blotting. ß-Actin serves as the loading control. **e.** The virus preparations from (**c**) and (**d**) were used to infect human organoids derived from fallopian tube. The organoids were selected for puromycin. The images show organoids under selection 2, 4 and 6 days respectively.

**Extended data Figure 4: L-tryptophan activates the pGSK3ß-c-Myc axis and rescues chlamydial infection.**

**a.** HeLa 229 cells were either left uninfected or infected with *Chlamydia (Ctr)* and treated with IFN-γ with or without L-tryptophan (W). After 30 hpi, the cells were analysed via Western blotting (n=3). **b.** HeLa 229 cells were infected with *Ctr* at MOI 1 for 24 h and or treated with IFN-γ and L-tryptophan (W). The cells were lysed, and the supernatant was used to infect freshly plated HeLa 229 cells and bacterial load was determined by Western blotting for cHSP60. **c.** HeLa 229 cells were either left untreated or were treated with different amounts of L-tryptophan (W). After 24 h the cells were lysed and analysed by Western blotting. **d.** HeLa 229 cells were infected with *Ctr* at MOI 1 for 30 h and treated with IFN-γ or indole. The cells were lysed and analyse by Western blotting for c-Myc and cHSP60 levels (n=3). **e.** The cells from (**d**) were lysed and the supernatant was used to infect freshly plated HeLa 229 cells to assess the infectivity of the progeny. **f.** HeLa 229 cells with AHT-inducible expression of shc-Myc cells were treated with 1 ng/mL AHT for 2 h. Cells were left untreated or were pre-treated for 2 h with 100 ng/mL L-tryptophan (W), infected with *Ctr* at MOI 1. The cells were lysed, the supernatant was used to infect fresh cells, and lysed 30 hpi to study infectivity. **g.** WII-U2OS cells with inducible expression of c-Myc were left untreated or treated with 100 ng/mL AHT and 10 ng/mL IFN-γ for 2 h. Cells were grown without L-tryptophan, and infected with *Ctr* at MOI 1 and lysed at 30 hpi for Western blot analysis. **h.** Cells from the experiment shown in (**g**) were used for infectivity assay. **i**. Cells from human fimbriae (human Fimb) were either left uninfected or infected with *Ctr* and treated with IFN-γ with or without L-tryptophan (W). The cells were analysed by Western blotting 24 hpi. cHSP60 shows the intensity of chlamydial infection and ß-Actin serves as loading control (n=3).

**Extended data Figure 5: Pathway analysis**

**a.-d.** WII-U2OS cells were either left uninfected or infected with *Ctr* at MOI 1 for 30 h. The infected cells were either left untreated or treated with just 10 ng/mL IFN-γ and 100 ng/mL AHT to induce expression of c-Myc. The cells were extracted, and metabolites were analysed by LC-MS. Quality controls and data normalization were performed and a principal component analysis (PCA) (**a**), and pathway analysis (control vs infected (**b**), infected vs infected IFN-γ treated (**c**), infected IFN-γ treated vs infected IFN-γ treated and c-Myc expressed (**d**)) are shown here. FDR < 0.05, Impact > 0.4.

**Extended data Figure 6: Urea cycle intermediates.**

**a.** Cartoon depicting TCA cycle and nucleotide biosynthesis in human cells. **b.** WII-U2OS cells were either left uninfected or infected with *Ctr* at MOI 1 for 30 h. The infected cells were either left untreated or treated with just 10 ng/mL IFN-γ and 100 ng/mL AHT to induce over expression of c-Myc. The cells were extracted, and metabolites were analysed by LC-MS. Data are presented as mean ± SD of triplicate wells. * indicates a p value < 0.05, ** indicates a p value < 0.01, *** indicates a p value < 0.001 and **** indicates a p value < 0.0001. The levels of aspartate, serine and glycine are shown. **c.** Cartoon depicting the flux of aspartate from the TCA cycle. Oxaloacetate from TCA is converted into aspartate by mitochondrial malate dehydrogenase. The glutamate-aspartate antiporter transports aspartate into the cytosol where it is shuttled into the urea cycle. **d-f.** WII-U2OS cells were either left uninfected or infected with *Ctr* at MOI 1 for 30 h. The infected cells were either left untreated or treated with just 10 ng/mL IFN-γ and 100 ng/mL AHT to induce over expression of c-Myc. The cells were extracted, and metabolites were analysed by LC-MS. Data are presented as mean ± SD of triplicate wells. * indicates a p value < 0.05, ** indicates a p value < 0.01, *** indicates a p value < 0.001 and **** indicates a p value < 0.0001. The levels of arginine, ornithine, citrulline, nucleotide triphosphate (ATP, GTP, CTP and UTP) (**e**), and nucleotide monophosphate (AMP, GMP, CMP and UMP) (**f**) are shown.

