## Supplementary figures and images for "c-Myc plays a key role in IFN-γ induced persistence of *Chlamydia trachomatis*"

### Fig. S1

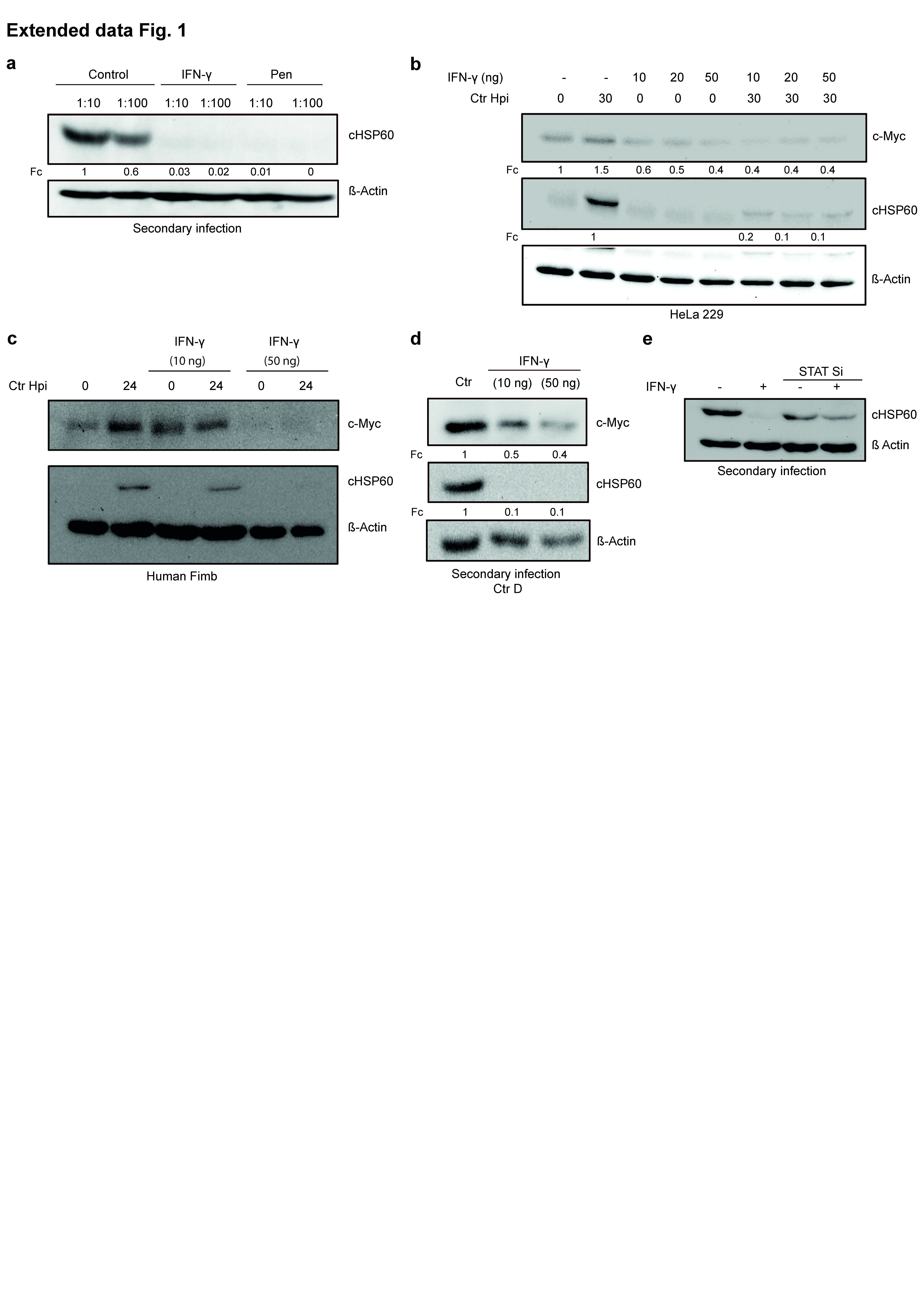

### Fig. S2

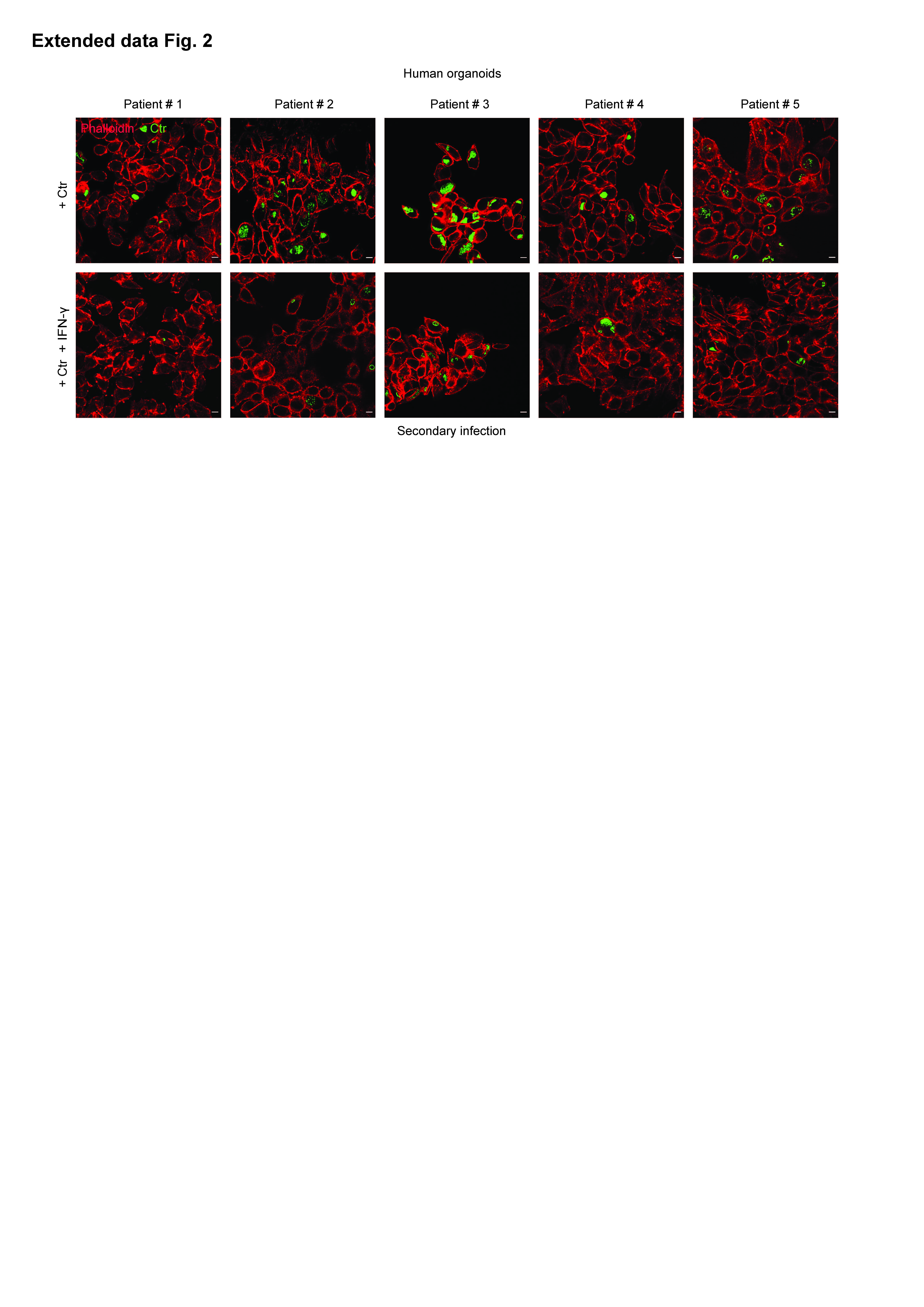

### Fig. S3

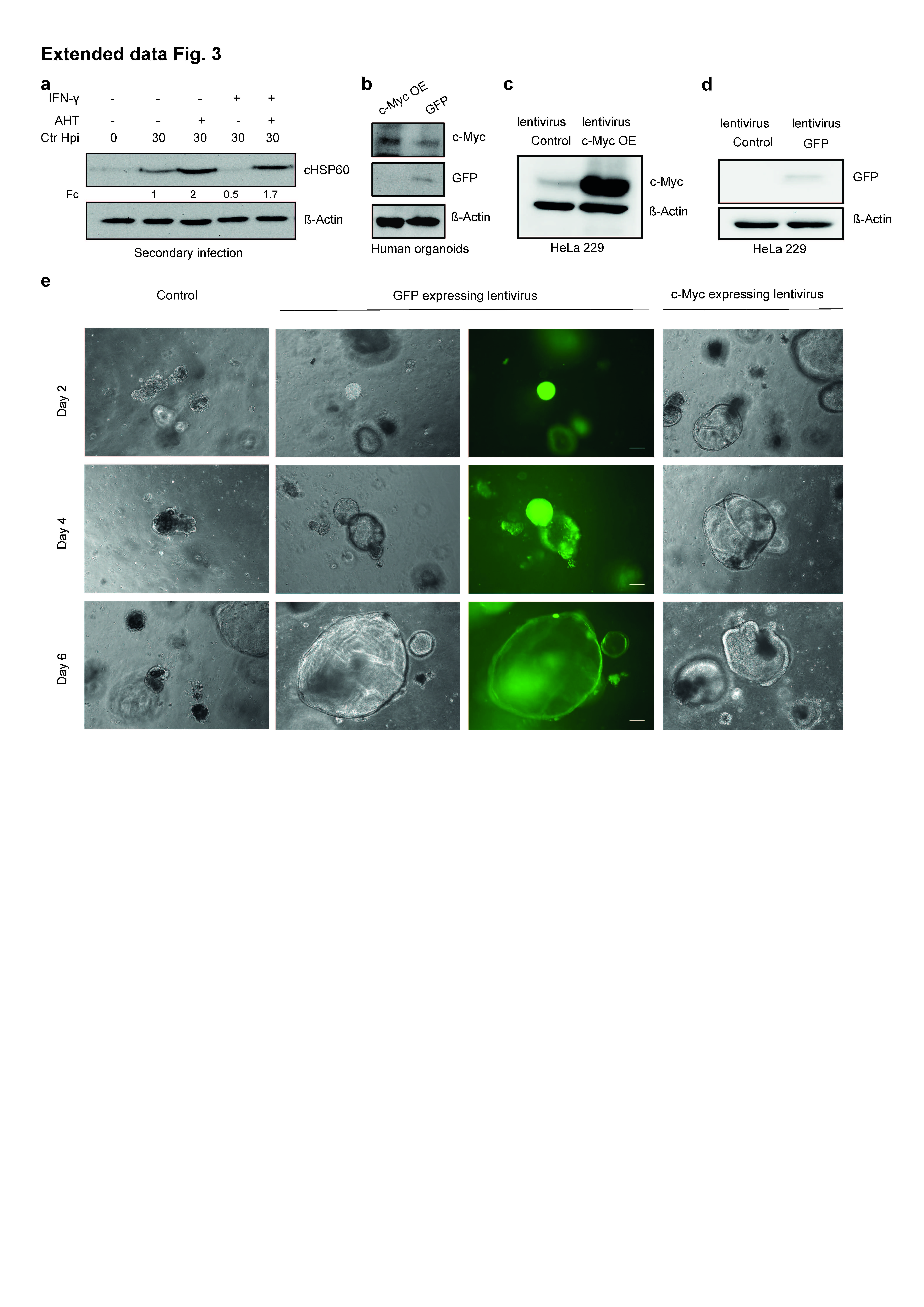

### Fig. S4

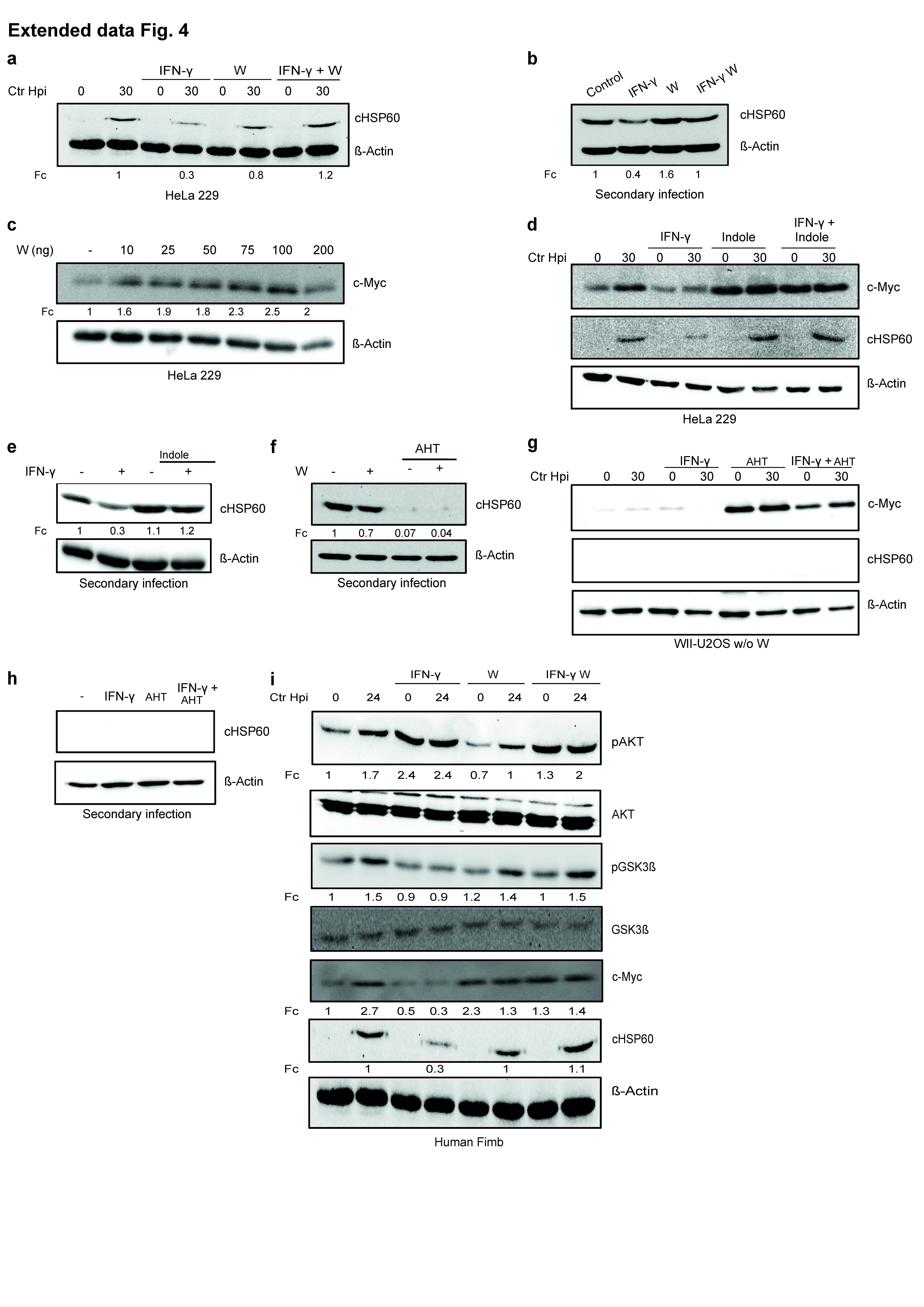

### Fig. S5

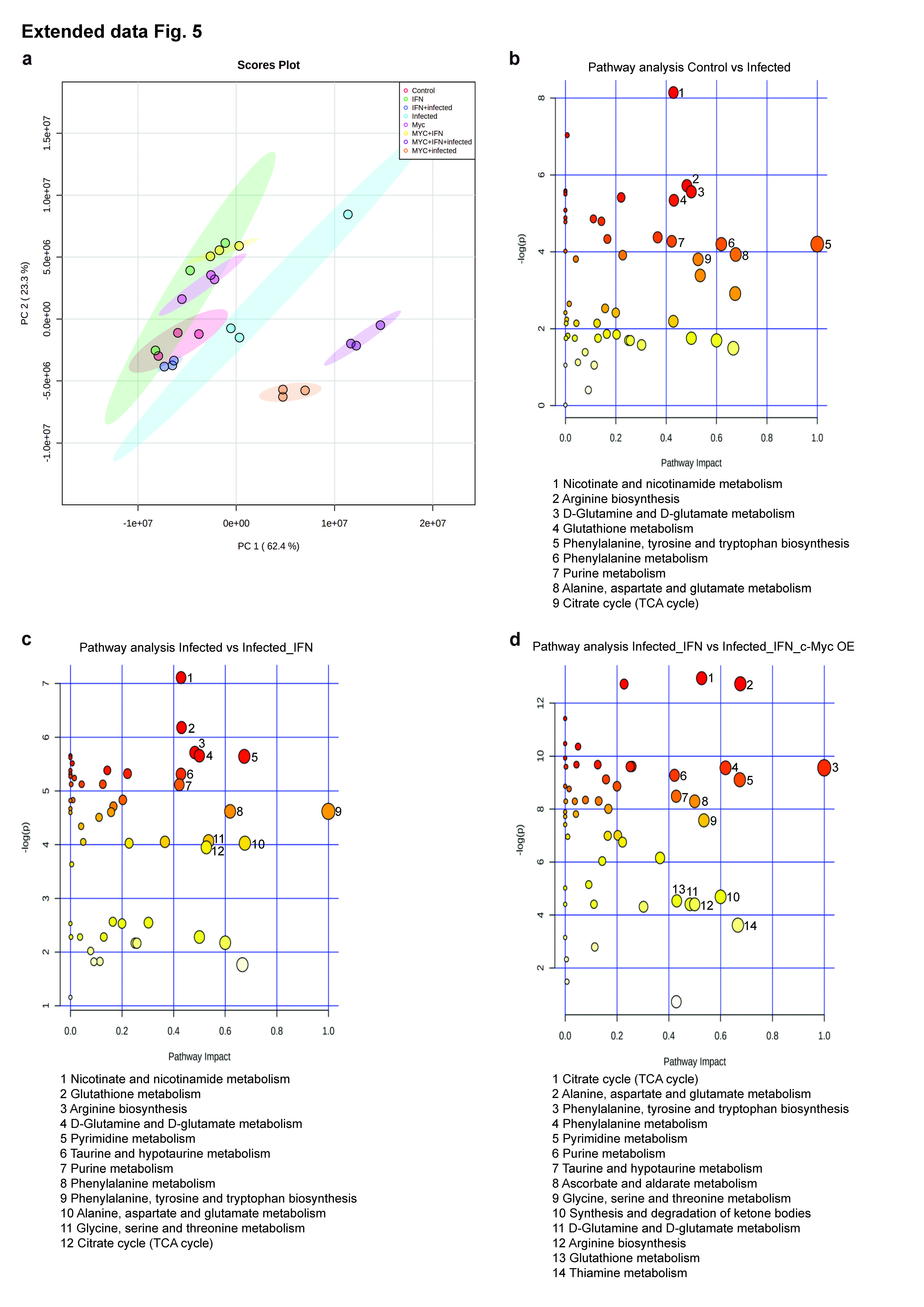

### Fig. S6

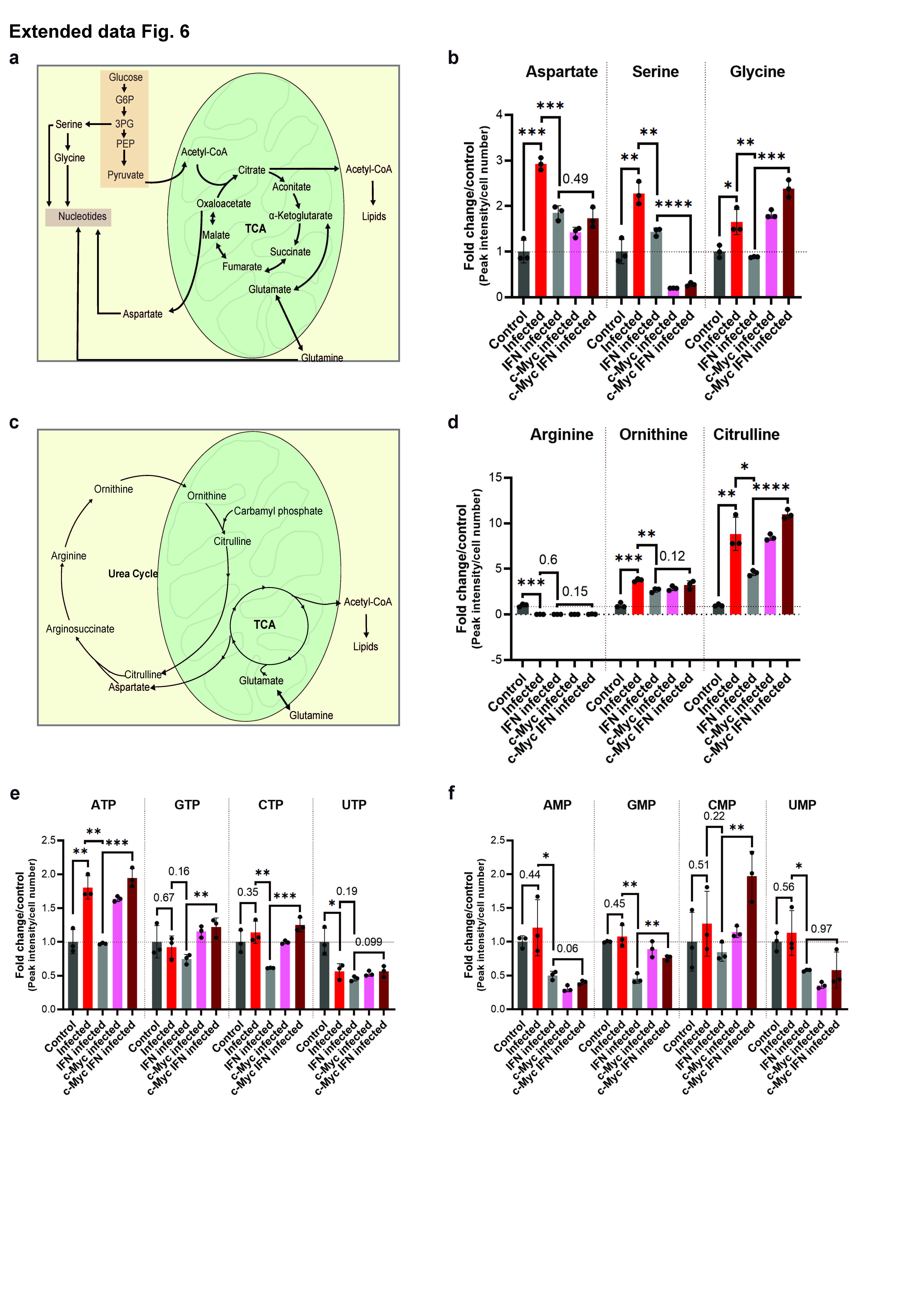
